# Refractive-index matching enhanced polarization sensitive optical coherence tomography quantification in human brain tissue

**DOI:** 10.1101/2021.09.12.459943

**Authors:** Chao J. Liu, William Ammon, Robert J. Jones, Jackson Nolan, Ruopeng Wang, Shuaibin Chang, Matthew P. Frosch, Anastasia Yendiki, David A. Boas, Caroline Magnain, Bruce Fischl, Hui Wang

## Abstract

The importance of polarization-sensitive optical coherence tomography (PS-OCT) has been increasingly recognized in human brain imaging. Despite the recent progress of PS-OCT in revealing white matter architecture and orientation, quantification of fine-scale fiber tracts in the human brain cortex has been a challenging problem, due to a low birefringence in the gray matter. In this study, we investigated the effect of refractive index matching by 2,2’-thiodiethanol (TDE) immersion on the improvement of PS-OCT measurements in *ex vivo* human brain tissue. We obtain the cortical fiber orientation maps in the gray matter, which reveals the radial fibers in the gyrus, the U-fibers along the sulcus, as well as distinct layers of fiber axes exhibiting laminar organization. Further analysis shows that index matching reduces the noise in axis orientation measurements by 56% and 39%, in white and gray matter, respectively. Index matching also enables precise measurements of apparent birefringence, which was underestimated in the white matter by 82% but overestimated in the gray matter by 16% prior to TDE immersion. Mathematical simulations show that the improvements are primarily attributed to the reduction in the tissue scattering coefficient, leading to an enhanced signal-to-noise ratio in deeper tissue regions, which could not be achieved by conventional noise reduction methods.

## 1. Introduction

Polarization-sensitive optical coherence tomography (PS-OCT) is an emerging technique to map large-scale *ex vivo* brain tissue [1-3]. In addition to the reflectivity which highlights the gross anatomy, PS-OCT utilizes the optical property of birefringence in the myelin sheath and provides quantitative image contrast metrics of myelinated axon bundles, including retardance and optic axis orientation. Nerve fiber tracts as small as tens of micrometers in diameter can be resolved from the retardance, and the orientation of nerve fibers can be measured by the optic axis orientation [4]. The use of PS-OCT to image intricate fiber architecture in the human brain has been demonstrated to be a source of ground-truth measurements of white-matter (WM) organization that does not suffer from the nonlinear distortions plaguing histological techniques [5-7]. One challenge of PS-OCT imaging in the human brain is to visualize the fibers in the gray matter (GM) and to quantify their axis orientation. While WM generates strong polarization-based signals, fiber tracts in the GM reveal significantly lower birefringence, and are thus difficult to visualize on retardance and orientation maps. Due to the strong scattering caused by tissue fixation, the OCT signal drops quickly as light propagates into the tissue, especially in regions deeper than 200 μm. This is especially a problem for PS-OCT systems using a single circular input state, in which the accuracy of retardance and axis orientation measurements strongly depends on signal-to-noise ratio (SNR). Low SNR can cause an inaccurate measurement of birefringence [8] and increase the noise in the axis orientation. In deeper regions where the SNR is low, the polarization measurements are unreliable and the estimation of apparent birefringence (Δ*n*) by fitting the slope of retardance in depth is also affected [9]. The limited SNR in PS-OCT images, especially in deeper regions affected by signal attenuation, compromises the validity of Δ*n* and axis orientation measurements. The improvement of accuracy in polarization measurements is crucial to map the human brain and other birefringent tissues using PS-OCT.

In recent years, refractive index matching media, such as glycerol [10], 2,2’-thiodiethanol (TDE) [11], and 3,3’-thiodipropanol [12], have shown the capability to increase the imaging depth for fluorescence imaging [13, 14]. The refractive index matching solution TDE can be miscible with water at any ratio and allows fine adjustment of the refractive index from that of water (1.33) to that of immersion oil (1.52). For OCT imaging, Yang et al. reported reduced scattering coefficient and improved imaging depth in *ex vivo* human brain via TDE immersion [15]. In this work, we investigate the effect of TDE immersion on polarization-based measurements of *ex vivo* human brain tissue using PS-OCT. We show significant improvement of optic axis orientation and birefringence estimations deep in the tissue from human brain samples, especially for the cortical fiber tracts in the GM. We then evaluate the improvements of index matching on SNR, birefringence and axis orientation at different depths and examine the effects at different TDE concentrations. Finally, we use simulation to explain the outcome of index matching on PS-OCT measurements by varying the noise, Δ*n*, and the attenuation coefficient (*α*) in a computational model. TDE immersion provides a promising approach to precisely measure polarization-based optical properties of human brain tissue using PS-OCT techniques.

## 2. Methods

## 2.1 Sample preparation

We obtained two brains from the Massachusetts General Hospital (MGH) Autopsy Suite for this study. Both subjects were neurologically normal prior to death (age at death: 69 and 60 years, 1 male and 1 female). The post-mortem interval did not exceed 24 hours. The brains were fixed by immersion in 10% formalin for at least two months and were cut into smaller blocks. We used a block from the occipital lobe of the first subject, and a block from the frontal lobe of the second subject. The tissue blocks were washed for one month in phosphate buffer saline solution (PBS) 0.01 M at room temperature while gently shaking. Refractive index matching was then performed with serial incubations in 20%, 40%, and 60% TDE in PBS (volume/volume) each for 24 hours at room temperature [15]. The *ex vivo* imaging procedures are approved by the Institutional Review Board of the MGH.

### 2.2 System setup and sample imaging

A home-made automatic serial sectioning PS-OCT (as-PSOCT) system [2] was used for data collection. The system integrated a commercial spectral domain PS-OCT centered at 1300 nm (TEL220PS, Thorlabs), motorized xyz translational stages, and a vibratome to section the tissue block. The imaging depth was 2.6 mm with an axial resolution of 4.2 μm in tissue. The maximum sensitivity of the system was 109 dB. Custom-built software, written in C++, coordinated data acquisition, xyz stage translation, and vibratome sectioning for automatic imaging of brain blocks. The samples were first flat faced with the vibratome and then imaged using a scan lens (OCT-LSM3, Thorlabs), yielding a lateral resolution of 10 μm. One volumetric acquisition composed of 400 A-lines and 400 B-lines covering a field of view of 2 mm × 2 mm took 8 seconds at an A-line rate of 50 kHz. Automatic tile-scan was used to cover the whole sample, with a 16% overlap between tiles. The samples were first imaged with PBS immersion before refractive index matching, and subsequently with 20%, 40%, and 60% TDE immersion after each incubation.

### 2.3 Image processing and quantitative analysis of PS-OCT metrics

The complex depth profiles of the PS-OCT data can be expressed in the form of *A*_H,V_(*z*) exp[*iϕ*_H,V_(*z*)], where *A* and *ϕ* denote the amplitude and phase as a function of depth *z*, and the subscripts H and V represent, respectively, the horizontal and vertical polarization channels. The image contrasts of reflectivity, *R*(*z*), retardance, *δ*(*z*), and optic axis orientation, *θ*(*z*) along depth, were obtained by *R*(*z*) ∝ *A*_H_(*z*)^2^ + *A*_V_(*z*)^2^, *δ*(*z*) = arctan[*A*_H_(*z*)/*A*_V_(*z*)] and *θ*(*z*) = [*ϕ*_H_(*z*) − *ϕ*_V_(*z*)]/2, respectively. To get the degree of polarization uniformity (DOPU), we first synthesized the Stokes vectors as [*I Q U V*]^T^ = [*A*_H_^2^ + *A*_V_^2^ *A*_H_^2^ − *A*_V_^2^ 2*A*_H_*A*_V_*cosθ* 2*A*_H_*A*_V_*sinθ*]^T^[16]. DOPU was then calculated as 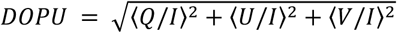, where ⟨ ⟩ denotes the local averaging operator within a 3-pixel Hanning shaped kernel. *En-face* retardance images were calculated by averaging the respective values along 500 μm in depth. *En-face* axis orientation images were formed by histogram analysis that relied on binning the orientation values along 500 μm into 5 deg intervals and calculating the mean value of a histogram-based Gaussian fit as the pixel value. The cumulative depth range of 500 μm was selected to reveal the cortical fibers in the GM due to the weak birefringence.

To examine the effect of index matching, we divided the A-line profiles into three depth ranges (0 - 200 μm, 200 - 400 μm, and 400 - 600 μm) and estimated the quantitative metrics of PS-OCT measurements in each segment. We quantified the piecewise attenuation coefficient *α* by fitting the slopes of the reflectivity profile (dB) in each depth interval. The piecewise SNR in reflectivity (*SNR*_*I*_) was calculated as *SNR* = 10. *log*_O_[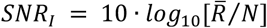/*N*], where 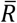 is the averaged reflectivity within a depth interval and *N* is the noise floor [8]. The apparent birefringence Δ*n* was calculated by performing a linear regression of the retardance profile and taking the absolute values of the slopes for every 200 μm interval. We also calculated *Var*_*θ*_, the standard deviation (s.t.d.) of optic axis orientation by fitting a Gaussian distribution to the axis orientation measurements in each of the three intervals. As gray and white matter carried significantly different birefringence and attenuation coefficients, we investigated the metrics above separately for each structure. In addition, we evaluated the effect of different TDE concentrations on those metrics.

### 2.4 Simulation of index matching effect on PS-OCT measurements

To simulate the PS-OCT contrasts, we first obtained the combined intensity *I*(*z*) from the two polarization channels along depth as *I*(*z*)= *I*_0_*exp*(−*αz*), where *I*_0_ is the intensity on tissue surface. Then the intensity of each polarization channel was obtained by *I*_H_(*z*) = *I*(*z*)*sin*^2^*φ*(*z*) + *N*_H_(*σ*_*n*_) and *I*_V_(*z*) = *I*(*z*)*cos*^2^*φ*(*z*) + *N*_V_(*σ*_*n*_) [17], where *φ*(*z*) = Δ*nz*. Additive noises following normal distribution with s.t.d. of *σ*_*n*_ were added to the two channels as *N*_H_ and *N*_V_, separately [18, 19]. Reflectivity and retardance with noise were calculated as 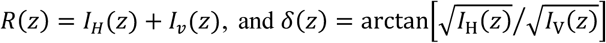. The phase in the two channels was obtained with the Hilbert transform and was then used to compute the *Var*_*θ*_ of axis orientation [20]. To investigate the effect of index matching, we varied *α* in the PS-OCT profile. We also inspected the influence of *σ*_*n*_ on the measurements. To simulate PS-OCT signals of gray and white matter, we evaluated the effects in low and high birefringence settings, respectively. We calculated Δ*n* from the slope of retardance and *Var*_*θ*_ from axis orientation in the 0 - 250 μm and 250 - 500 μm depth interval, following the definition described in section 2.3. We simulated 1000 A-lines and obtained the average Δ*n* and *Var*_*θ*_ in the two depth intervals.

## 3. Results

### 3.1 Refractive index matching increased PS-OCT contrasts in human brain tissue

We first investigated the effects of 60% TDE on human primary visual cortex in the occipital lobe. TDE immersion increased the effective imaging depth remarkably in cross-sectional reflectivity, retardance and axis orientation images (right panels in Fig. 1a and Fig. 1b). We observed banded patterns in cross-sectional retardance and orientation of the WM indicating the presence of high birefringence (dashed rectangular boxes in the right panel of Fig. 1b), whereas the patterns were completely undermined by the noise in deep regions without index matching (right panel of Fig. 1a). In addition to the much-improved imaging quality in WM, we also clearly visualized the cortical fiber tracts in the GM in cross-sectional retardance and orientation images (white arrows in the right panel of Fig. 1b), which were suppressed by the noise with PBS immersion (right panel of Fig. 1a).

**Fig. 1.**
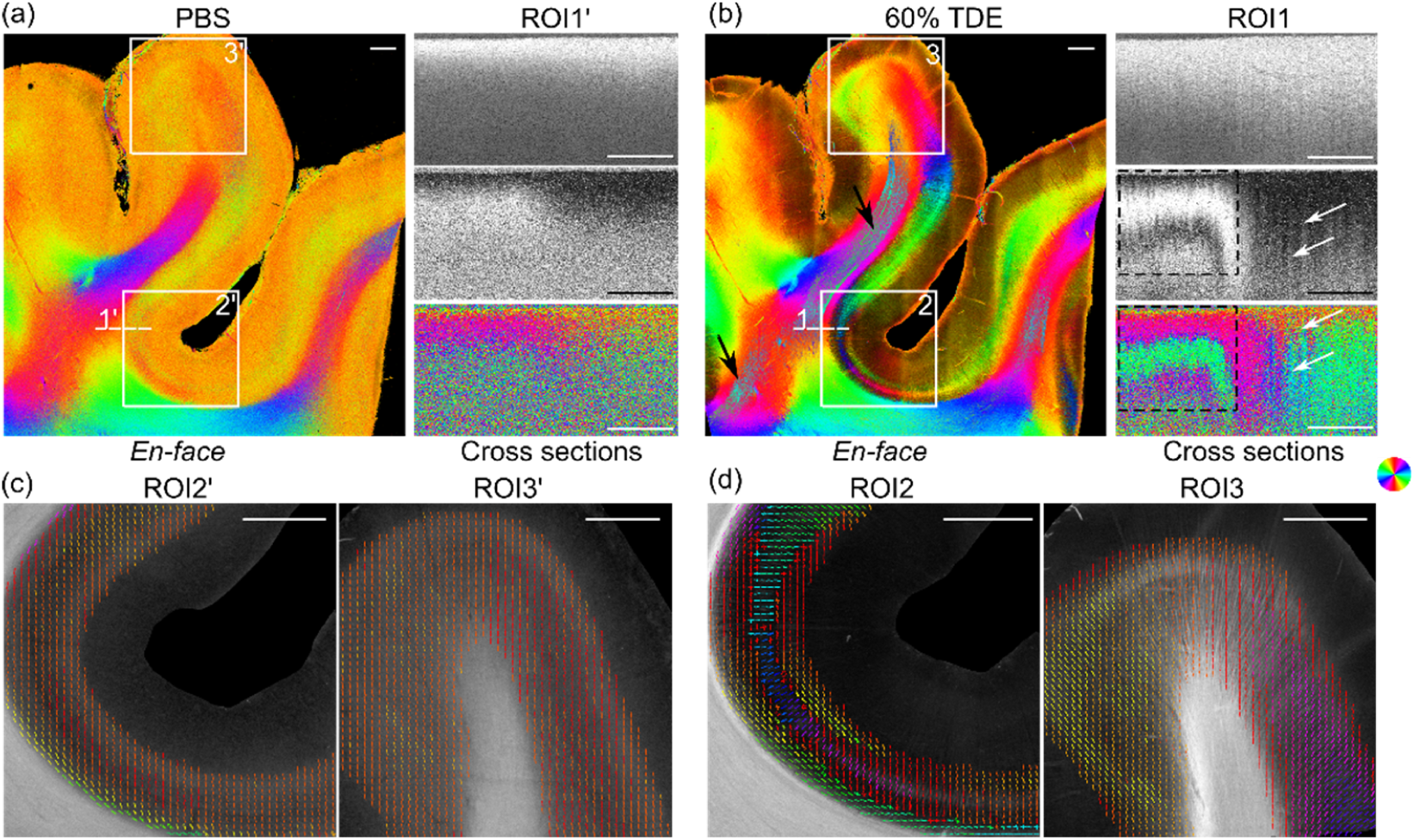
PS-OCT images of human visual cortex in the occipital lobe *ex vivo* with PBS (a, c) and 60% TDE immersion (b, d). (a)-(b) *En-face* optic axis orientation images are shown on the left and cross-sectional images are shown on the right. The location of the cross-sectional images of reflectivity (35∼75 dB), retardance (0∼60 deg) and optic axis orientation (−90∼90 deg) were indicated by the dashed line (1’ and 1) on the *en-face* orientation in (a) and (b), separately. Black arrows in the *en-face* orientation of (b) indicate the artifacts in deep WM. Dashed rectangular boxes on the cross-sectional image of (b) highlight the banded pattern in retardance and axis orientation in the WM. White arrows in the cross section of (b) indicate cortical fiber tracts in the retardance and axis orientation images in the GM. (c)-(d) *En-face* retardance images superimposed with fiber polar maps show enlarged region in the sulcal area (ROI2’ and ROI2) as well as gyral area (ROI3’ and ROI3). The polar plots are derived from the *en-face* orientation. For orientation images and polar plots, axis orientation values are color coded in HSV space as illustrated by the color wheel and the brightness of orientation images is modulated by retardance. Scale bars: 1 mm.

We generated the *en-face* axis orientation images with PBS and 60% TDE immersion using the 0 - 500 μm depth range. With 60% TDE, the axis orientation map in the sulcus revealed the stripe of Gennari, containing tangential fibers in layer IV of the primary visual cortex, as well as an additional stripe directly below the stripe of Gennari, characterized by radially oriented fibers spanning several layers (ROI2 in Fig. 1b). In the gyral area, the radial fibers can be seen across different layers (ROI3 in Fig. 1b). In a striking contrast, these cortical fiber tracts were indistinguishable when imaged with PBS immersion. It is noticeable that some artifacts in the highly birefringent WM area are present in the *en-face* orientation image with 60% TDE immersion (black arrows in the left panel of Fig. 1b). Since we used 500 μm depth range for *en-face* orientation formation, the banding pattern in the cross-sectional orientation caused these artifacts (see Section 2.3 in Methods and dashed rectangular boxes in right panel of Fig. 1b).

To further compare the quality of orientation estimation in cortical fiber tracts, we plotted the polar maps of two ROIs with PBS and 60% TDE immersion (Fig. 1c and 1d). The *en-face* orientation image was divided into patches of 80 μm × 80 μm areas, in which the fiber orientation distribution functions were constructed by generating histograms of orientation measurements in each patch, using a bin width of 5 deg. The polar plots were displayed as the histograms. We only showed layers 4-6 to highlight the more birefringent region in the cortex. In the sulcus region (ROI2), the polar maps from TDE immersion allowed us to clearly distinguish the stripe of Gennari and the radial fibers. Another stipe of cortical fibers in layers 5/6 (so called U-fibers), whose orientation aligned with the white matter, was also revealed (Fig. 1d). In the gyrus (ROI3), the enhanced imaging quality with TDE immersion revealed the fanning geometry of cortical fibers (Fig. 1d). This was not possible with PBS, due to the rapid drop of SNR in deep regions. In that case, the polar map was dominated by noise and most of the polar plots pointed to 90/-90 deg (Fig. 1c).

We also imaged a tissue section from the frontal lobe of another subject and observed a similar effect by index matching (Fig. 2). In addition to the increased SNR in orientation maps, we can accurately measure the orientation of radial fibers (see lower panel in Fig. 2) across different layers in the cortex as well as the U-fibers in deep cortex (see white arrows in Fig. 2) connecting adjacent gyri. Those cortical fibers cannot be revealed by measurements with PBS immersion.

**Fig. 2.**
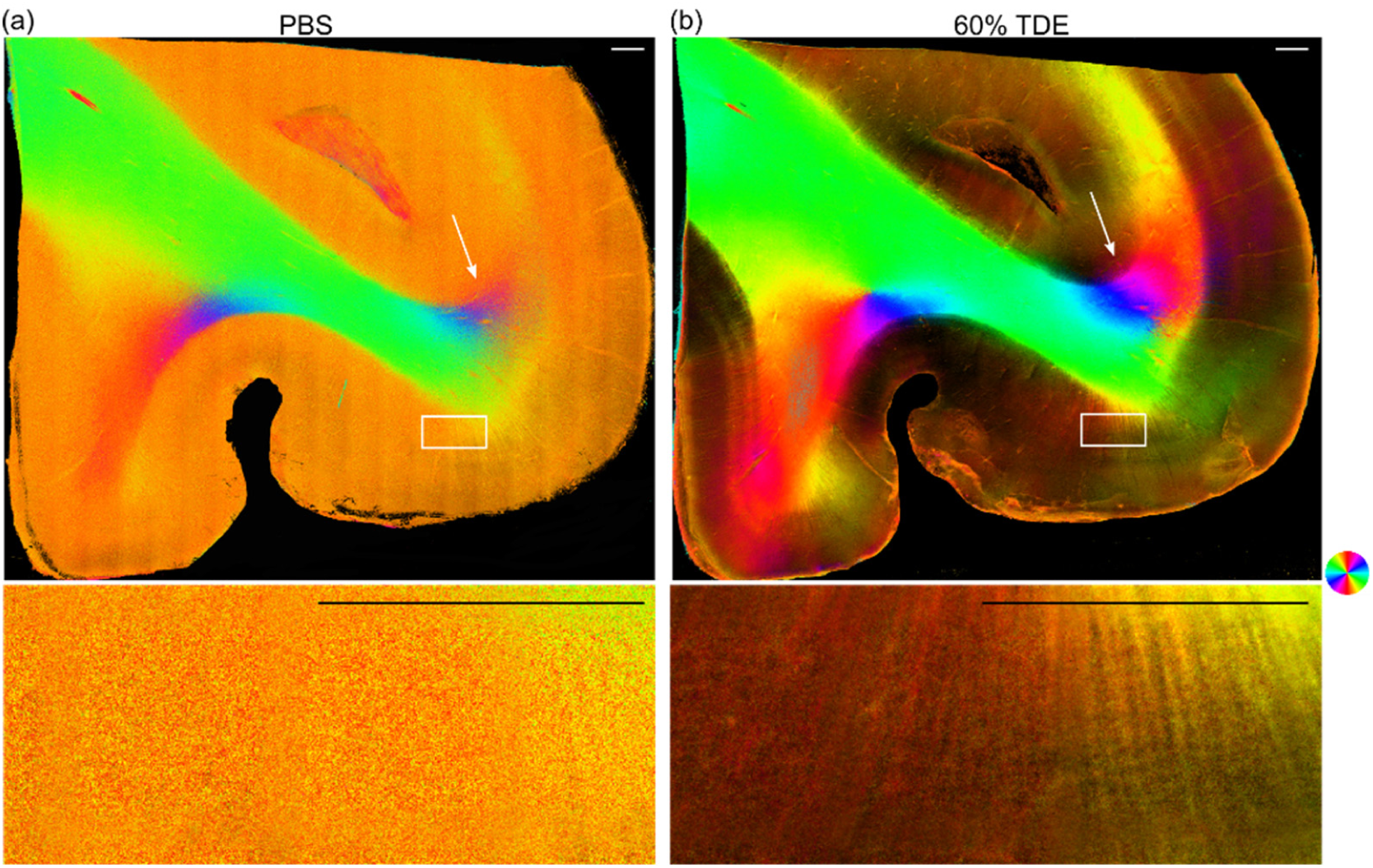
*En-face* axis orientation images of human frontal lobe *ex vivo* from PS-OCT with PBS (a) and 60% TDE immersion (b). White arrows indicate the U-fiber. Rectangular boxes indicate an area of radial fibers enlarged in the lower panel. The orientation is defined by the color wheel and the brightness is modulated by retardance. Scale bars: 0.5 mm.

### 3.2 Refractive index matching improved the quantitative estimation of PS-OCT measurements

To evaluate the estimation of optical properties, we chose a 2 mm × 2 mm area from the occipital lobe sample (centered on ROI1’/ROI1 in Fig. 1a and Fig. 1b). We manually segmented the WM and GM using the retardance image for the analysis in Fig. 3-Fig. 4. We examined the depth profiles of reflectivity, retardance, axis orientation, and DOPU in GM and WM, respectively, and compared the results measured with PBS and 60% TDE immersion.

**Fig. 3.**
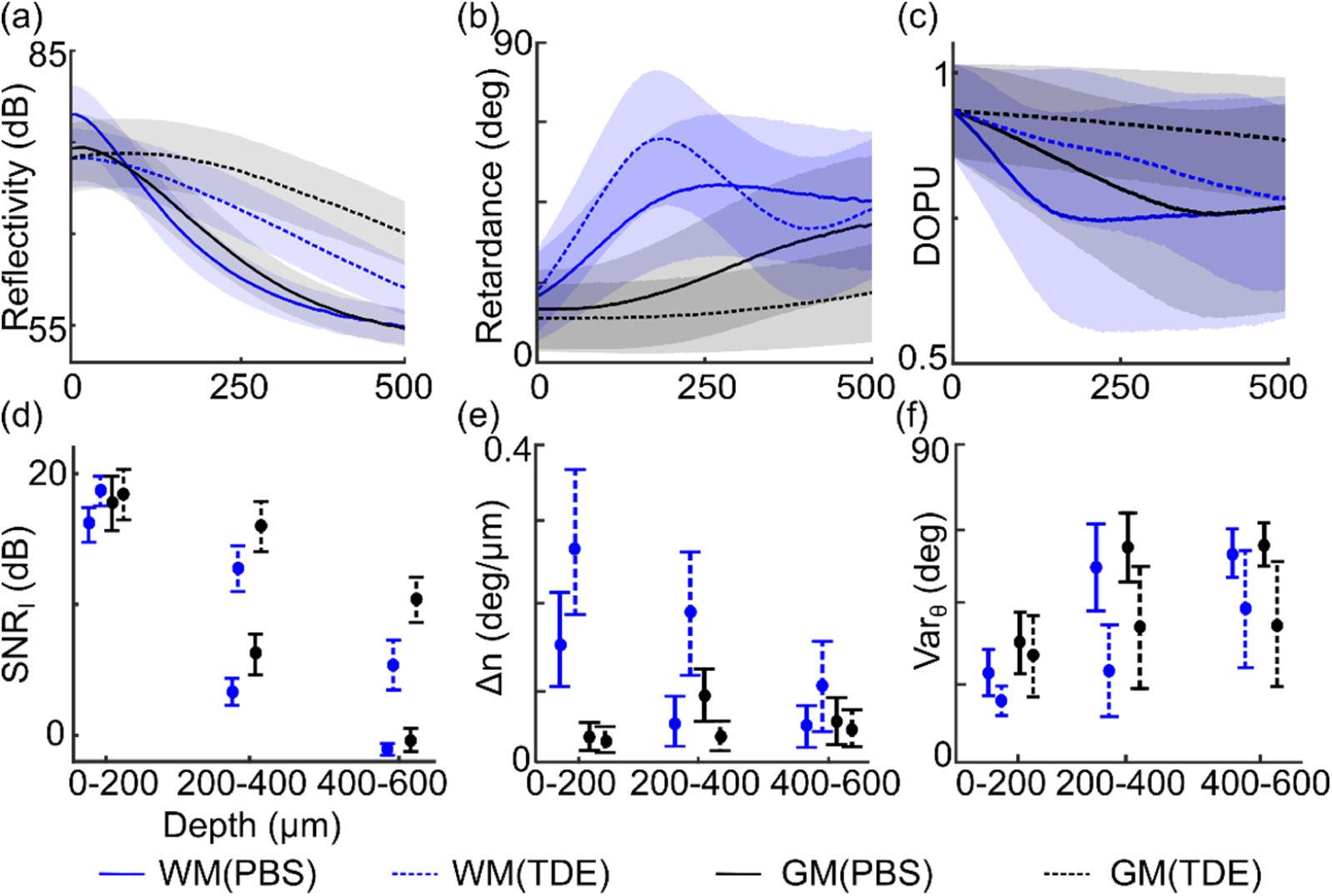
TDE immersion improves the SNR and enhances PS-OCT quantification. (a)-(c) Reflectivity (a), retardance (b) and DOPU (c) along depth for WM and GM with PBS and 60% TDE immersion. The shaded areas define the s.t.d. of A-lines. (d)-(f) *SNR*_*I*_ (d), Δ*n* (e) and *Var*_*θ*_ (f) calculated from three depth ranges with PBS and 60% TDE immersion, respectively. The error bars indicate s.t.d. of A-lines and the mean values are shown as dots. In all panels, blue and black curves indicate WM and GM, respectively; solid and dashed lines indicate measurements with PBS and 60% TDE immersion, respectively.

**Fig. 4.**
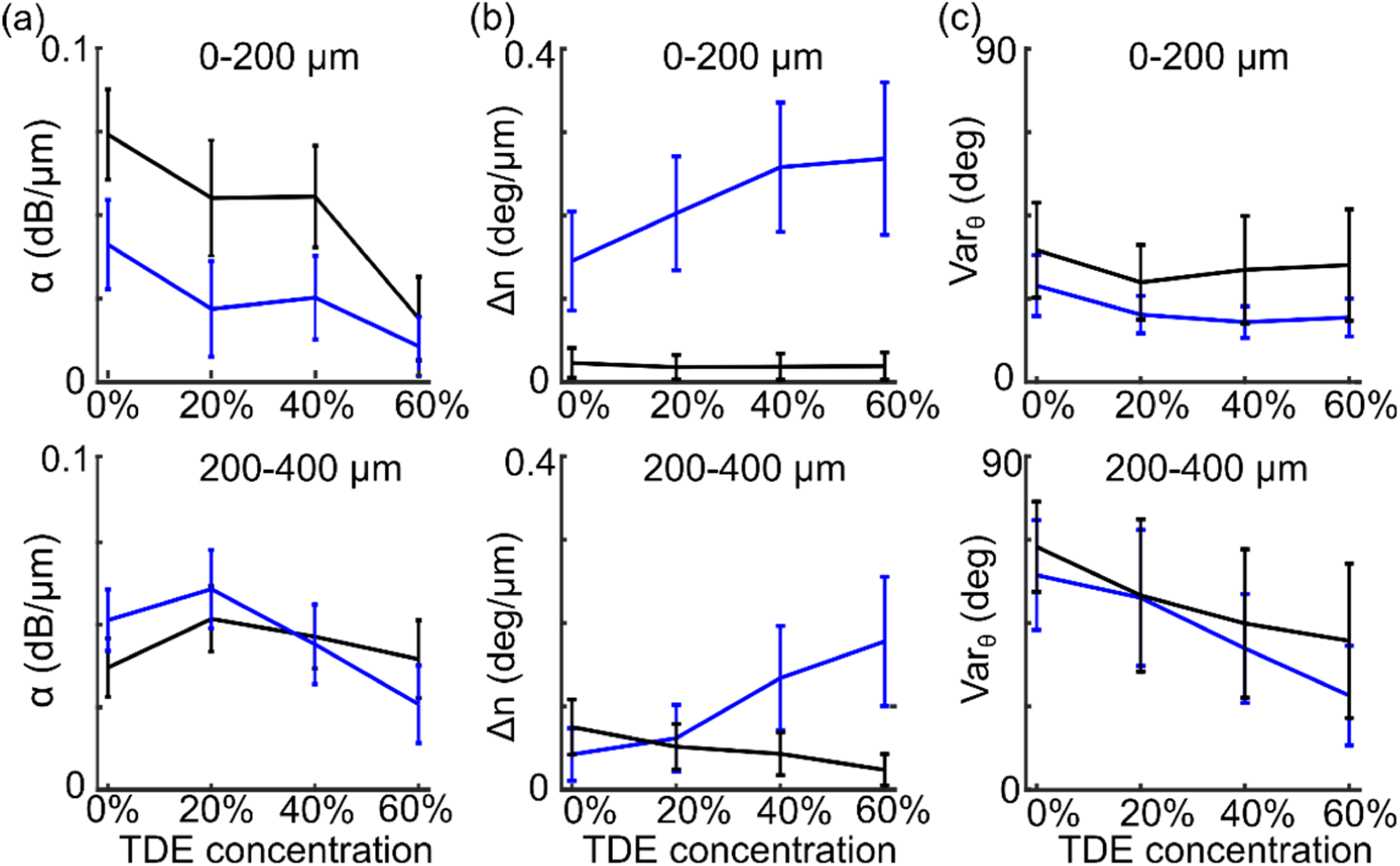
PS-OCT measurements with respect to TDE concentration. Attenuation coefficient *α* (a), apparent birefringence Δ*n* (b) and s.t.d. of axis orientation *Var*_*θ*_ (c) using 0-200 μm (top panel) and 200-400 μm (bottom panel) interval at 0%, 20%, 40% and 60% TDE concentration. The error bars indicate s.t.d. of A-lines, and the blue and black curves indicate WM and GM, respectively.

For reflectivity, 60% TDE immersion reduced light attenuation both in GM and WM compared to PBS immersion (Fig. 3a) [15]. We examined the *SNR* of reflectivity at different depth ranges (Fig. 3d). In the first 200 μm, the *SNR*_*I*_ of GM (black) and WM (blue) was similar between PBS (solid line) and 60% TDE (dashed line) immersion. The greatest improvement occurred in the deeper region where signal dropped quickly with PBS immersion. In the range 200-400 μm, *SNR*_*I*_ with 60% TDE immersion was improved by 9.4 dB and 9.8 dB relative to PBS in WM and GM respectively. In the 400-600 μm range, *SNR*_*I*_ dropped close to 0 dB both in WM and GM with PBS immersion, indicating no distinction between signal and noise floor (55 dB). However, with 60% TDE immersion, the *SNR*_*I*_ was still at 6.5 dB and 10.8 dB above the noise floor in WM and GM, respectively. TDE immersion greatly increased the SNR, compared to PBS immersion.

We then examined the retardance profiles from which apparent birefringence can be estimated. With 60% TDE immersion, we observed a significantly improved delineation of retardance measurements both in WM and GM. In the WM, with 60% TDE immersion the retardance cumulation caused by high birefringence remained visible up to 500 μm deep, which peaked at 195 μm and formed a banded pattern in 200-400 μm range and a third band at depths beyond 400 μm (Fig. 3b, dashed blue line). In contrast, with PBS immersion the retardance profile became flat after 250 μm where the amplitude of both polarization channels fell to a similar noise floor, i.e., *A*_H_ ≅ *A*_V_ (Fig. 3b, solid blue line). In the low birefringent GM, with 60% TDE immersion retardance remained low as expected along the 500 μm depth range, only elevated by 12 deg (Fig. 3b, black lines). In contrast, with PBS immersion retardance measure was erroneously elevated in depths greater than 200 μm and reached ∼40 deg at 500 μm due to the diminished SNR in both polarization channels. The improved retardance profiles resulted from TDE immersion would contribute to a more accurate estimation of Δ*n*.

The improvement of Δ*n* was elaborated at different depth intervals (Fig. 3e). In the WM (blue bars), the superficial depth range (0-200 μm) yielded a Δ*n* of 0.27±0.09 deg/μm with 60% TDE immersion, which was 82% higher than that with PBS immersion. The difference was more remarkable in deeper regions. Δ*n* was calculated to be 0.18±0.08 deg/μm at the 200-400 μm interval and 0.09±0.06 deg/μm at the 400-600 μm interval with 60% TDE immersion; whereas Δ*n* was unreliable at both intervals with PBS immersion as retardance fell into the noise level. The low SNR with PBS immersion caused a noticeably lower estimation of Δ*n* in WM, especially when fitting the slope of retardance in the deeper (and hence noisier) regions. Conversely, Δ*n* in GM revealed an opposite trend. The 0-200 μm regime showed a 16% lower estimation with 60% TDE immersion compared to PBS (black bars in Fig. 3e, 0.019±0.02 deg/μm with 60% TDE immersion versus 0.023±0.02 deg/μm with PBS immersion). In the 200-400 μm range, Δ*n* stayed low with 60% TDE immersion; however, noise heavily influenced both polarization channels with PBS immersion and Δ*n* was elevated by 68%. Overall, 60% TDE immersion enabled a much higher birefringence estimation in WM while a substantially lower estimation in GM, and the difference between TDE and PBS immersions was more pronounced in deeper and nosier regions.

We also calculated the DOPU from experimental data with PBS and 60% TDE immersion. DOPU characterized the depolarization properties of the tissue [21]. The DOPU decayed fast along depth with PBS immersion (solid lines in Fig. 3c), especially in WM and thus the polarization states were scrambled. With 60% TDE immersion, the delay of DOPU became slower in both GM and WM indicating more preserved polarization states in tissue (dashed lines in Fig. 3c).

For the orientation measurements, the values of WM were reliable with PBS immersion in the first 200 μm (Fig 1a), yielding a low *Var*_*θ*_ of 26 deg. However, with 60% TDE immersion we still observed a decreased *Var*_*θ*_ by 33% and 11%, respectively, in WM and GM (Fig. 3f). The difference is more pronounced in deeper and noisier regions. In the 200-400 μm range, the *Var*_*θ*_ in both WM and GM became much higher in PBS due to low SNR (*Var*_*θ*_= 58±15 deg and 65±12 deg, respectively); 60% TDE immersion greatly reduced *Var*_*θ*_ by 56% and 39% to 25±13 deg and 40±21 deg in WM and GM, respectively. The effect of noise suppression in orientation with TDE immersion preserved in the even noisier 400-600 μm range, where *Var*_*θ*_ with 60% TDE immersion was 26% and 35% lower in WM and GM, respectively, compared to that with PBS immersion. TDE immersion greatly reduced the *Var*_*θ*_ and hence increased the reliability in axis orientation measurement.

We further evaluated the effect of TDE concentration on PS-OCT measurements. We quantified the attenuation coefficient, the birefringence, and the *Var*_*θ*_ of orientation with 0%, 20%, 40%, and 60% TDE immersion at two depth ranges: 0-200 μm and 200-400 μm (Fig. 4). The attenuation coefficient *α* in GM and WM both deceased with respect to TDE concertation (Fig. 4a). We observed a 69% decrease (0.084 ±0.009 dB/μm to 0.026±0.009 dB/μm) in the WM and an 86% decrease (0.054±0.011 dB/μm to 0.008±0.005 dB/μm) in the GM, from PBS to 60% TDE immersion within 0-200 μm range. *α* kept dropping with 40% and 60% TDE immersion in the deeper 200-400 μm range. In birefringence estimation, we found that the fitted Δ*n* increased in the WM but decreased in the GM with TDE concentration, and this trend was more pronounced in the deeper region. In the superficial WM, we measured Δ*n* was 39% higher with 20% TDE immersion than PBS immersion, which was further increased to 0.25±0.06 deg/μm with 40% TDE immersion and remained stable with 60% TDE immersion (Fig. 4b upper panel, blue lines). In the 200-400 μm range, the trend of fitted Δ*n* increase with respect to TDE concentration remained (46%, 217% and 321% higher, respectively, in 20%, 40% and 60% TDE than that with PBS immersion, Fig. 4b lower panel, blue lines). In the GM, fitted Δ*n* was similarly low in 20%, 40% and 60% TDE in the 0-200 μm range (mean value: 0.018 deg/μm). In the 200-400 μm, we observed a more pronounced effect of TDE concentration, where Δ*n* estimation became erroneously high with PBS immersion (0.075±0.04 deg/μm) and was lowered by 31%, 43% and 68% with respect to 20%, 40%, and 60% TDE immersion, respectively (Fig. 4b, black lines). Finally, the reliability of orientation measurement improved with the increase of TDE concentration, which manifested a substantial decrease in *Var*_*θ*_ at the deeper regions (200-400 μm, Fig. 4c). *Var*_*θ*_ was not greatly affected by TDE concentration in the 0-200 μm range as the SNR remained high at superficial depth in all of the solutions. In the 200-400 μm interval, *Var*_*θ*_ of WM reduced by 10%, 34% and 56%, respectively, with 20%, 40% and 60% TDE immersion than PBS immersion, and *Var*_*θ*_ of GM decreased by 20%, 32% and 39%, respectively.

### 3.3 Simulation of refractive index matching effects on PS-OCT measurements

To understand the effects of refractive index matching on polarization-based quantification by PS-OCT, we simulated PS-OCT measurements to evaluate the contribution of two factors to the accuracy of the quantitative metrics (Δ*n* and *Var*_*θ*_), including the attenuation coefficient and the noise. Separate simulations were performed for the WM and GM, in which birefringence was significantly different.

We simulated the effects of refractive index matching by varying *α* with given Δ*n* (high birefringence of 0.27 deg/μm for WM and low birefringence 0.02 deg/μm for GM, respectively, in Fig. 5a and Fig. 5b). The standard deviation *σ*_*n*_ of noise was set to be 8 dB, which was comparable with our experimental data. For WM, the banded pattern of the retardance depth-profiles was only present with low *α*, where SNR remained high in the deeper region (the left panel in Fig. 5a). By calculating the slope of retardance, we found that the estimated Δ*n* was very close to set value (0.27 deg/μm) at *α* = 0.01 dB/ μm in both 0-250 μm and 250-500 μm depth ranges. However, the estimated Δ*n* was negatively correlated with *α* (green lines in Fig. 5a), and this negative correlation was more dramatic in the noisier 250-500 μm depth range (the right panel in Fig. 5a). The simulation results indicated that Δ*n* in WM was significantly underestimated without refractive index matching such as with PBS immersion. Conversely, for the low birefringent GM, consistent with experimental observation, retardance failed to remain low but was quickly elevated to 45 deg noise level with high attenuation coefficient (see *α* = 0.1 dB/μm in Fig. 5b). By reducing *α*, the increasing trend of retardance over depth became slower (see *α* = 0.01 dB/μm in Fig. 5b). Consequently, Δ*n* estimation was accurate at low *α* (see *α* = 0.01 dB/μm in Fig. 5b); however, estimated Δ*n* was positively correlated with *α* while in effective SNR regime (green lines in Fig. 5b). The simulation indicated that high attenuation without refractive index matching caused substantial overestimation of Δ*n* in GM as observed in our experimental data. In 250-500 μm range, the estimated Δ*n* became lower again for *α*>0.04 dB/μm (see green line in right panel of Fig. 5b), because the retardance signal became flat at the noise level. Overall, the simulation in Fig. 5a and Fig. 5b explained our experimental observations: Δ*n* was underestimated in WM but overestimated in GM without refractive index matching due to the high attenuation.

**Fig. 5.**
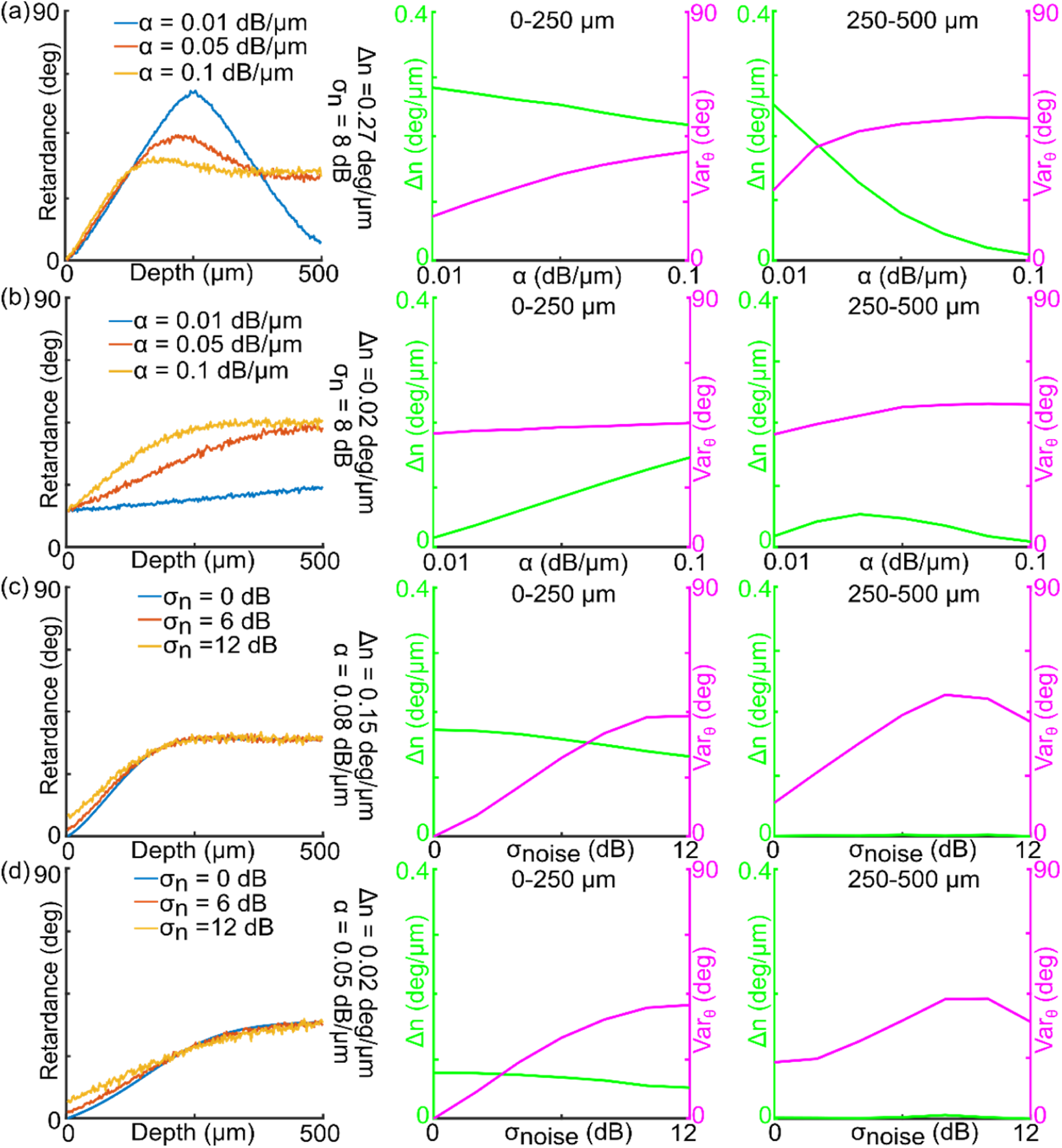
Simulation of the effects of attenuation coefficient *α* (a, b) and noise level *σ*_*n*_ (c, d) on PS-OCT quantifications in WM (a, c) and GM (b, d) of human brain tissues. (a) – (d) Simulated retardance along depth (left panel), the calculated Δn from retardance and Var_θ_ from axis orientation in the 0-250 μm interval (middle panel) and 250-500 μm interval (right panel). The parameters used in the simulations are labeled in each panel. In (a) and (b), *α* is varied with given *σ*_*n*_ and Δ*n*. In (c) and (d), *σ*_*n*_ is varied with given *α* and Δ*n*. Green and magenta lines indicate the calculated Δn and Var_θ_, respectively.

The *Var*_*θ*_in axis orientation measurement was also affected by *α*. In WM, *Var*_*θ*_ increased with *α* and the deeper 250-500 μm region had higher *Var*_*θ*_ compared to the superficial depths (magenta lines in Fig. 5a). For GM, *Var*_*θ*_ was higher compared to WM due to low birefringence and consequently low signal in the cross-polarized channel (magenta lines in Fig. 5b). Reducing attenuation coefficient effectively decreased *Var*_*θ*_ both in WM and GM and hence increased the reliability of orientation measurement.

In addition, we simulated the effect of different noise levels with fixed attenuation coefficients, for both WM (*α* = 0.08 dB/μm, Δ*n* = 0.15 deg/μm, Fig. 5c) and GM (*α* = 0.05 dB/μm, Δ*n* = 0.02 deg/μm, Fig. 5d). The parameters were set to match the experimental measurements without refractive index matching. We found that Δ*n* of both WM and GM in the superficial depth was slightly underestimated as the noise increased from 0 dB to 12 dB (the middle panel in Fig. 5c and 5d), which indicated that reducing noise can only slightly correct Δ*n* underestimation at the superficial depth. However, in the deeper range, retardance approached and remained at 45 deg in both GM and WM regardless of the noise level, and thus the Δ*n* estimation by fitting the slope of retardance failed (the right panel in Fig. 5c and Fig. 5d). As expected, higher noise also caused greater *Var*_*θ*_ in orientation. Taking the attenuation and the noise effects together, our simulation elaborated that the improved PS-OCT quantification via refractive index matching was primarily attributed to the reduced light attenuation, which could not be achieved by purely reducing the noise in each polarization channel.

## 4. Discussion

Neurons in the brain communicated via axonal projections, which originate and terminate in the gray matter and create connectivity via fiber bundles in the white matter. Despite the recent progress of PS-OCT in imaging white matter fiber architecture and orientation, the quantification of complex, fine-scale fiber tracts in the cortex have not been reported. In this work, we described the first demonstration of human cortical fiber orientation maps with PS-OCT. We reported the distinctive, laminar organization of the fiber axis in human visual cortex, as well as the fiber orientation patterns in gyri and sulci in both occipital and frontal cortex samples. Moreover, we identified the orientation of U-fibers at the GM-WM border, which connect adjacent gyri in the human brain. The clinical implications of abnormalities in U-fiber regions have been investigated by recent diffusion weighted magnetic resonance imaging (dMRI) studies in various neurological disorders, such as multiple sclerosis [22] and Alzheimer’s disease [23]. In this work, we showed that PS-OCT with TDE immersion can characterize the microscopic properties of intracortical connections. In the future, it can be applied to study how these properties change in neurodegenerative disease. In addition, an important application of PS-OCT is to obtain ground truth axonal orientations for the validation of dMRI in WM [6, 7]. Similarly, PS-OCT with TDE immersion can be applied to validate dMRI measurements in the GM, which are challenging due to low anisotropy and limited resolution [24, 25].

The orientation of cortical fiber tracts has typically not been the focus in previous PS-OCT studies [2, 4]. This is because the low birefringence in the GM does not allow sufficient signal cumulation along depth on the cross-polarized channel before intensity decays to the noise floor due to high tissue scattering. Consequently, the phase measurement enabling optic axis orientation is not reliable within the effective imaging range. Instead, the orientation measurement is biased by an optic axis caused by a non-negligible system birefringence when compared to the GM birefringence. The problem is especially prominent in PS-OCT systems using a single circular input state. As seen in the current study, cross-sectional orientation image without refractive index matching was dominated by one color and lack of fiber orientation distinction in the superficial depth, while it was masked by noise in deeper regions. This led to the universal yellow-orange appearance on *en-face* orientation across the GM region without TDE immersion (Fig. 1a & Fig. 2a). After refractive index matching, we observed a remarkable improvement in cortical fiber orientation distinction (Fig. 1b & Fig. 2b). Due to the increased SNR, TDE immersion substantially reduced the noise in axis orientation measurements, especially in the 200-400 μm depth range (by 56% and 39% in WM and GM, respectively, Fig. 3f). The improvement of the orientation measurement in deeper regions makes it possible to increase the effective imaging depth of PS-OCT up to 400 μm in the *ex vivo* human brain.

In traditional PS-OCT imaging, quantification of cortical fiber birefringence has been challenging due to substantial speckle noise and limited SNR. It has been shown that improving SNR can increase the accuracy of polarization-based measurements in PS-OCT [9]. To this end, numerous noise reduction methods have been employed in PS-OCT [8, 19, 26, 27]. Here our simulation results showed a limited improvement of birefringence quantification by denoising (Fig. 5c and 5d). The high attenuation coefficient in *ex vivo* tissues quickly deteriorated the OCT signals along depth making the retardance measurement unreliable. In this work, we proposed using refractive index matching to reduce the attenuation rate and hence significantly increased the SNR in the deeper region. Our data showed substantially improved accuracy for estimating Δ*n* in both GM and WM, which was verified by simulation results (Fig. 5a and 5b). Particularly, we found that without refractive index matching, the apparent birefringence was underestimated by 82% in WM while overestimated by 16% in GM even at the superficial depth range (Fig. 3e). The estimation of birefringence at depths greater than 200 μm without refractive index was completely unreliable. This was further elaborated by the simulation in Fig. 5a and Fig. 5b, where retardance gradually approached 45 deg, regardless of high or low birefringence due to the loss of SNR. By using refractive index matching via TDE immersion, we achieved a substantial enhancement of SNR along PS-OCT depth profiles. As a result, the birefringence estimation demonstrated a remarkable improvement in both superficial and deep regions of highly scattered brain tissue.

TDE allows a precise setting of the refractive index (n) by adjusting the concentration and here we imaged the brain sample with a TDE concentration from 0% (n = 1.33) to 60% (n = 1.44) [11]. Previous work showed that TDE reduced scattering in human brain tissue in a concentration-dependent manner [15]. Here we found the increase of TDE concentration improved the accuracy of the apparent birefringence (Fig. 4b) and reduced the noise in the axis orientation (Fig. 4c) in the superficial depth range with up to 20%-40% TDE immersion; however, both measurements continue to improve in the deeper 200-400 μm range with 60% TDE immersion (Fig. 4a and 4c). The refractive index of GM and WM has been reported to be 1.367 and 1.467, respectively [15]. Future work will include the investigation of optimal volume concentration of TDE for maximizing image features and contrast of interest in PS-OCT.

The improvement in imaging depth via TDE immersion has a significant benefit for as-PSOCT in reconstructing large-volume three-dimensional (3D) human brain tissues, in which sequential cutting of 100 μm thin slices was typically required for water immersed samples [2]. Our study suggests that refractive index matching has the potential to reduce number of slicing by a factor of 4 due to increased effective imaging depths. Therefore, our work paves the way to further improve data collection efficiency for large-scale human brain mapping. One interesting direction of this work is to develop depth-resolved fiber orientation mapping algorithms [28-30]. Current orientation information is extracted as *en-face* images assuming a constant fiber axis along imaging depth, which is problematic for refractive index matching enhanced PS-OCT, as extensive fiber crossing can be present within 400 μm depth range in human brain. As a result, local birefringence and orientation retrieval is imperative in volumetric human brain imaging in order to achieve microscopic fiber orientation mapping [31, 32].

Besides PS-OCT, three-dimensional polarized light imaging (3D-PLI) provides quantitative analysis of myelinated axons in thin brain sections based on rotating polarimetry [33]. The transmission mode measurements using incoherent light eliminated the influence of speckle noise on the microscopic images. The axis orientations of cortical fibers have been quantified by 3D-PLI and have been utilized to better understand brain connectivity such as the transcallosal connection of the visual cortex [34]. Despite the accurate quantification within a slice, the registration of 3D-PLI images between consecutive sections is still difficult, due to the nonlinear distortions induced by sectioning and tissue mounting [35]. In contrast, the block-face imaging procedure of as-PSOCT obviates the need for between-slice registration in volumetric reconstruction. Therefore, as-PSOCT aided by refractive index matching is a promising tool to reconstruct the 3D cortical fiber orientation and myeloarchitecture in large-scale human brain samples at micrometer accuracy.

The improvement of optic axis orientation measurements can also greatly benefit the estimation of through-plane fiber axis by PS-OCT using variable illumination angles [36]. Conventional PS-OCT based on one illumination angle only measures the in-plane fiber angle. Estimating the full 3D orientation requires also measuring the through-plane fiber angle. Moreover, the precise measurement of birefringence provides a solid foundation for estimating the true birefringence which can be retrieved by apparent birefringence and through-plane fiber angle [37].

## 5. Conclusions

In summary, we investigated the effect of refractive index matching for PS-OCT measurements of *ex vivo* human brain tissue. TDE immersion increased the SNR in deeper regions due to reduced attenuation. We visualized and quantified the orientation of cortical fibers in GM, including the Stria of Gennari, radial fibers and U-fibers. The noise of orientation measurements reduced by 56% and 39% in WM and GM in deeper regions, where orientation values were typically unreliable without refractive index matching. The improvement of the SNR enabled precise estimation of apparent birefringence by fitting the slope of retardance profile along depth. We found apparent birefringence measured without refractive index matching could be underestimated by 82% in WM while overestimated by 16% in GM even at superficial depths. TDE immersion provides a promising approach to precisely measure the optical properties of human brain tissue using PS-OCT. The increased imaging depth and SNR could pave the road to a comprehensive mapping of fiber architecture in the human brain using as-PSOCT.

## Data availability

Data underlying the results presented in this paper are not publicly available at this time but may be obtained from the authors upon reasonable request.

